# Single-cell transcriptome analysis reveals estrogen signaling augments the mitochondrial folate pathway to coordinately fuel purine and polyamine synthesis in breast cancer cells

**DOI:** 10.1101/246363

**Authors:** Detu Zhu, Xianglan Zhaozu, Guimei Cui, Shiehong Chang, Yi Xiang See, Michelle Gek Liang Lim, Dajiang Guo, Xin Chen, Paul Robson, Yumei Luo, Edwin Cheung

## Abstract

Estrogen regulates diverse physiological effects and drives breast tumor progression by directly activating estrogen receptor α (ERα). However, due to the stochastic nature of gene transcription and the resulting heterogeneous cellular response, it is important to investigate estrogen-stimulated gene expression profiles at the single-cell level in order to fully understand how ERα regulates transcription in breast cancer cells. In this study, we performed single-cell transcriptome analysis on ERα-positive breast cancer cell lines following 17β-estradiol stimulation. Overall, we observed robust gene expression diversity between individual cells. Moreover, we found over two thirds of the genes in breast cancer cells displayed a bimodal expression pattern, which caused averaging artifacts and masked the identification of potential estrogen-regulated genes. We overcame this issue by reconstructing a dynamic estrogen-responsive transcriptional network from discrete time points into a pseudotemporal continuum. Pathway analysis of the differentially expressed genes derived from the pseudotemporal analysis showed an estrogen-stimulated metabolic switch that favored biosynthesis and cell proliferation but reduced estrogen degradation. In addition, we identified folate-mediated one-carbon metabolism as a novel estrogen-regulated pathway in breast cancer cells. Notably, estrogen stimulation reprogramed this pathway through the mitochondrial folate pathway to coordinately fuel polyamine and de novo purine synthesis. Finally, we showed AZIN1 and PPAT, key regulators in the above pathways, are direct ERα target genes and essential for breast cancer cell survival and growth. In summary, our single-cell study illustrated a dynamic transcriptional heterogeneity in ERα-positive breast cancer cells in response to estrogen stimulation and uncovered a novel mechanism of an estrogen-mediated metabolic switch.

## Introduction

Breast cancer is the most prevalent cancer in women worldwide with more than two-thirds diagnosed as estrogen receptor α (ERα)-positive^1^. Estrogen is a steroid hormone that plays pivotal functions in the pathophysiology of ERα-positive breast cancers^2^. Specifically, estrogen binds to ERα which subsequently is recruited to estrogen response elements (EREs) in the genome. ERα then recruits a host of cofactor proteins to either activate or repress the transcription of target genes that are essential for tumor proliferation and progression^3^.

Global time-series gene expression studies have revealed a list of ERα-regulated genes and the underlying mechanisms of estrogen-mediated cancer development^4^. However, the initiation of ERα binding to chromatin is stochastic^5^. Moreover, an individual cell’s response to stimuli is further complicated by a variety of intrinsic and extrinsic factors such as chromatin status, mitochondrial content, relative location within a population, and cell-to-cell contact^6^. Hence, the classical time-series gene expression analysis based on bulk cell population averages may be incomplete due to the multi-layered stochasticity in cellular responses to estrogen stimulation. Thus, new technologies to analyze transcriptomes at the single-cell level are required to account for these variables.

Recent advances in live-cell imaging have enabled researchers to monitor the dynamic expression of target genes in individual cells and confirmed that gene expression at the single-cell level is highly stochastic and binary^7^. However, these techniques are limited by the number of reporters and detection channels, and are unable to facilitate de novo discovery^8^. The availability of automated microfluidic platforms that can uniformly capture and process hundreds of single-cells for RNA-seq analysis has enabled profiling of single-cell transcriptomes at discrete time points^9^. Even so, gene expression cannot be tracked continuously from the same single-cell, and the stochasticity of response to stimulation will exist across the time-series^10^. Nevertheless, by leveraging on new computational algorithms, we can construct a pseudotemporal continuum through the reorganization of single-cell transcriptomes from discrete time points^11^. In these trajectory analyses, the transcriptome from each single-cell represents an instance or a “pseudotime point” along an artificial time vector that denotes the progress along the stimulus response^12^. Thus, by constructing a pseudotemporal continuum we can obtain an enhanced temporal resolution compared to analyzing each discrete experimental time point.

In this work, we performed time-course studies with 17β-estradiol (E2) treatment followed by single-cell RNA-sequencing (scRNA-seq) analysis on MCF-7 and T47D ERα-positive human breast cancer cell lines. We applied “Monocle” single-cell trajectory analysis and ordered cells along an artificial pseudotemporal continuum to allow the characterization of the unsynchronized response of breast cancer cells to estrogen stimulation. With an increased temporal resolution, we uncovered a new mechanism of an estrogen-mediated metabolic switch and identified novel ERα-regulated genes that are essential for these reprogrammed metabolic pathways.

## Results

### Single-cell RNA-seq analysis of breast cancer cells in response to estrogen stimulation

To characterize the extent of gene expression variability in individual breast cancer cells in response to estrogen stimulation, we performed single-cell RNA-seq analysis on ERα-positive MCF-7 and T47D cells harvested at 0, 3, 6 and 12 hours after E2 stimulation (Figure 1A). We captured a total of 177 (85 MCF-7 and 92 T47D) single-cells and sequenced each cell to an averaged depth of 10 million reads (Supplementary Table 1). After filtering for unique mapping reads (Supplementary Figure 1) and 3’ or 5’ coverage biases (Supplementary Figure 2), we retained 84 MCF-7 and 78 T47D single-cell profiles (Supplementary Table 1) for further downstream analysis. We then estimated the expression level of all the UCSC-annotated genes and removed genes that were not appreciably expressed (FPKM>1 in more than half of the single-cell profiles). In the end, 6,867 genes were used for our analysis.

**Figure 1.**
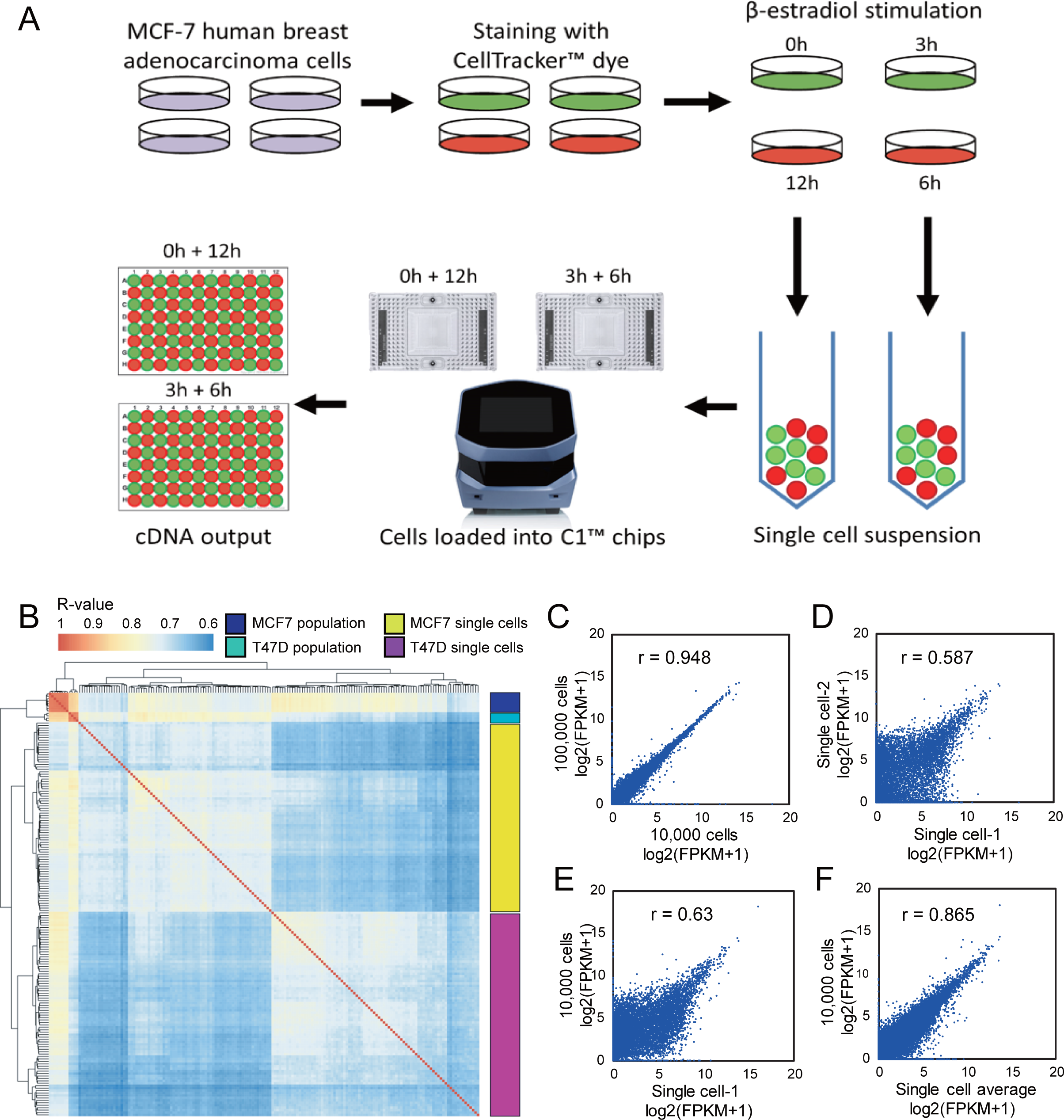
Generation of scRNA-seq data from MCF-7 and T47D cells under estrogen stimulation. **(A)** Workflow depicting the rapid and high-throughput isolation and RNA-seq of single-cells from breast cancer cell lines under time-course E2 treatment. **(B)** Correlation between single-cell profiles and bulk population profiles. **(C)** Scatter plot of gene expression values (log2(FPKM+1)) between the 100k cell bulk profile and the 10k cell bulk profile (r=0.948). **(D)** Scatter plot between 2 random single-cell profiles with low correlation (r=0.587). **(E)** Scatter plot between the 10k cell bulk profile and a random single-cell profile showing low correlation (r=0.63). **(F)** Scatter plot between the 10k cell bulk profile and the average of all single-cell profile (r=0.865).

Next, we performed Pearson correlation coefficient (PCC) analysis followed by unsupervised clustering to measure the linear correlation between all the single-cell samples and the population replicates from the two cell lines. As shown in Figure 1B, MCF-7 and T47D cells are clearly separated, indicating that cell line differences are greater than cell-to-cell variances. Although the gene expression level of the bulk population replicates closely correlated with one another (Pearson r > 0.94; Figure 1B and C), substantial differences in expression between individual cells exist (0.65 < r <0.8; Figure 1B and D; Supplementary Figure 3). Despite this extensive cell-to-cell variation, the averaged expression level of the single-cells correlated well with the bulk population replicates (Figure 1E and F).

To validate the inter-cellular heterogeneity observations from our scRNA-seq analysis, we performed droplet digital PCR (ddPCR), a high-resolution approach for quantifying the absolute number of mRNA copies in a tiny amount of sample. We assayed for the expression level of TFF1, GREB1, and ESR1 (three well-known E2-responsive marker genes), as well as ACTB (a housekeeping gene) (Supplementary Figure 5 and 6). Our ddPCR results showed there is a large cell-to-cell variance for every gene, including the housekeeping gene (Supplementary Figure 7), however, the overall expression profile of the 3 E2-responsive genes correlated well with the scRNA-seq results (Figure 2A and B; Supplementary Figure 8). Besides ddPCR, we also used single molecule RNA-fluorescence in situ hybridization (smRNA-FISH), an amplification-free imaging technique, to examine the expression of the above marker genes but in a greater number of cells. Similar to ddPCR, results from the smRNA-FISH analysis also mirrored the expression trends of the scRNA-seq data for both the MCF-7 (Figure 2C and D) and T47D (Supplementary Figure 9) cell lines. Taken together, our results showed that breast cancer cells displayed great transcriptional heterogeneity in response to estrogen stimulation.

**Figure 2.**
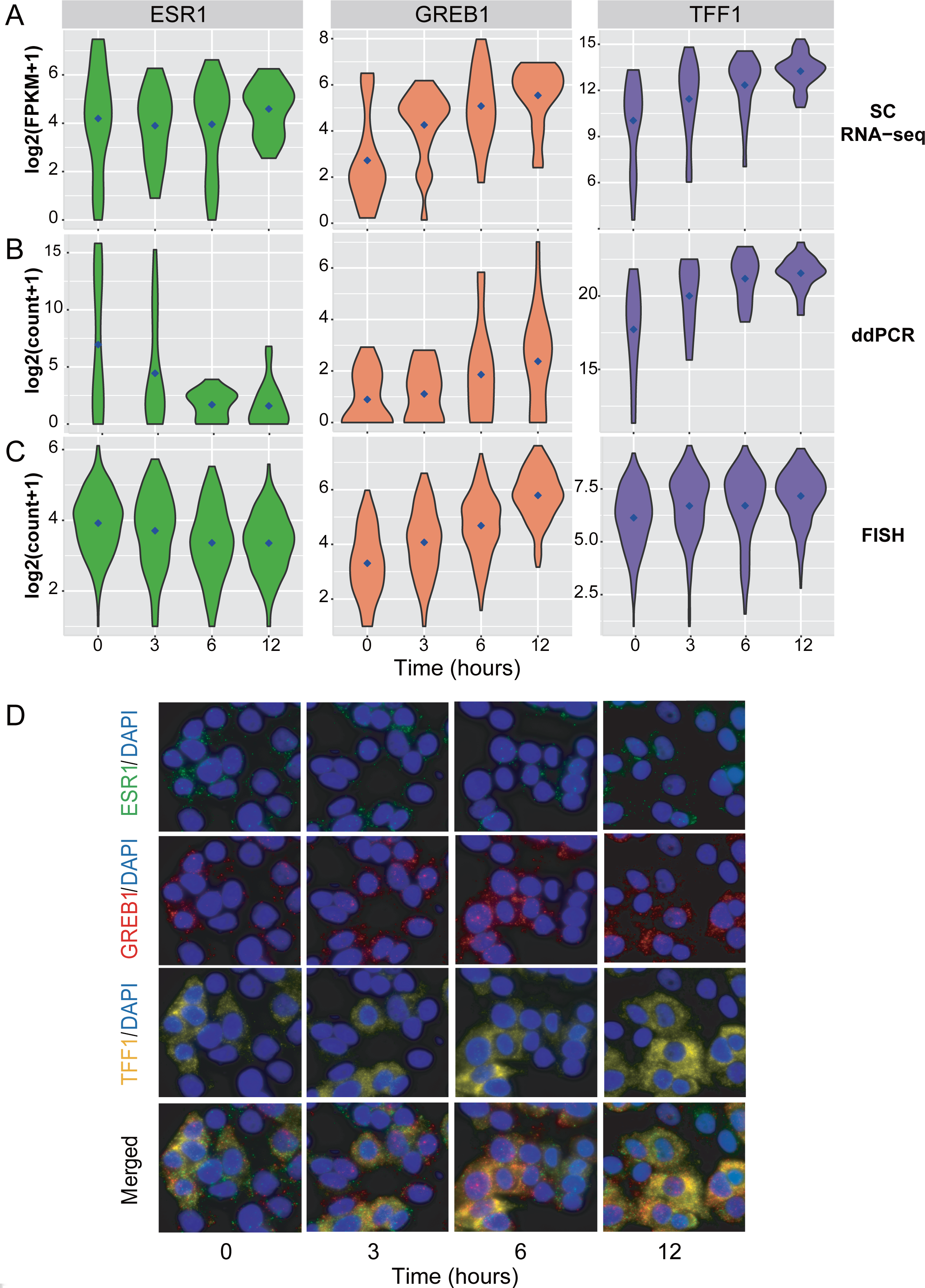
Validation of estrogen-responsive marker genes by smRNA-FISH and ddPCR. **(A)** Violin plots showing the scRNA-seq expression values (log_2_(FPKM+1)) of estrogen-responsive marker genes TFF1, GREB1, and ESR1 from the time-course E2 treatment of MCF-7 cells. **(B)** Violin plots showing single-cell ddPCR values (log_2_(Count+1)) of the marker genes from the same MCF-7 time-course experiment. **(C)** Violin plots showing smRNA-FISH values (log_2_(Count+1)) of the marker genes in a time-course E2 treatment of MCF-7 cells. 200 cells were counted for each time point. **(D)** Multiplexed smRNA-FISH images showing the mRNA copies of the marker genes in MCF-7 cells under time-course E2 treatment with nuclei counter stained by DAPI.

### Bimodal gene expression is common across single-cells and produces averaging artifacts

The mRNA expression level of genes frequently operate in two modes: baseline and over- or under-expression, which is a major contributor of cell-to-cell variances^9^. Thus, to characterize the extent of bimodal gene expression in our breast cancer scRNA-seq data, we used a recently developed algorithm called SIBER^13^. SIBER works by fitting the single-cell gene expression distribution into 2 log-normal distributions (component 1 and 2) and then calculating the mean values (Mu1 and Mu2) and the ratios (Pi1 and Pi2) for these two subpopulations (Figure 3A). Finally, a bimodal index (BI) value is calculated for each gene. Using SIBER, we calculated the BI value for every coding gene across all the single-cell profiles at each time point and determined any gene with a BI>1.8 as a bimodal gene (Figure 3B). As shown in Figure 3C, a large proportion (27-41%) of the genes in both MCF-7 and T47D cells are bimodal in each time point. Moreover, the BI values of each gene changed across time points. For example, approximately 6-8% of genes were constantly bimodal across the 4 time points, while >60% of the genes altered between biomodality and unimodality (Figure 3D).

**Figure 3.**
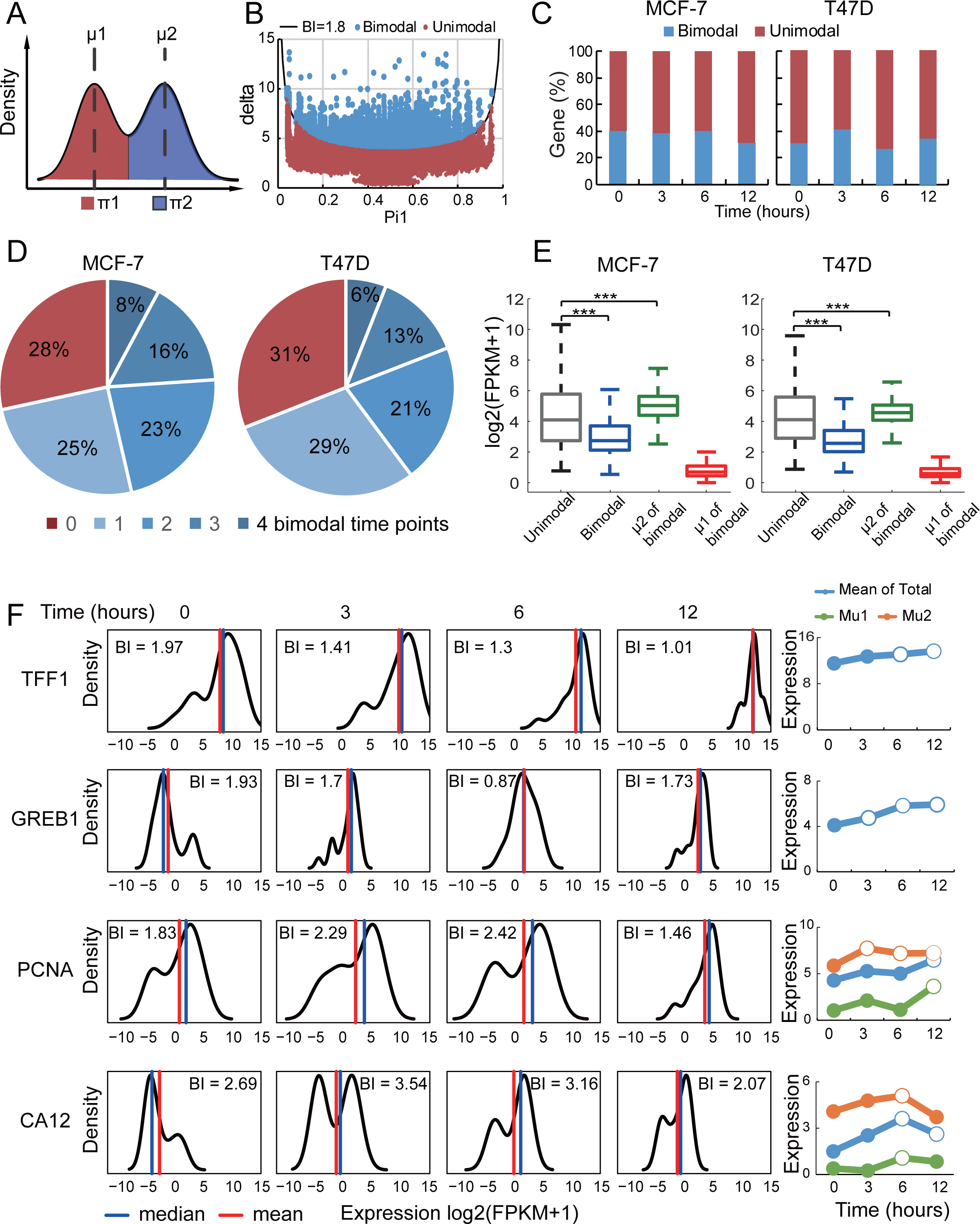
Bimodal variation of gene expression levels across single-cells causes averaging artifacts. **(A)** The Bimodality Index (BI) of each gene across single-cells in every time point was calculated by the SIBER method which is based on the ratio and distance between two components. Pi1 is the ratio of the lower component, Pi2 is the ratio of the higher component; Mu1 is the mean of the lower component, Mu2 is the mean of the higher component. **(B)** Genes with BI&1.8 (blue dots above the black curve) are defined as bimodal expression. **(C)** Percentage of genes with bimodal expression in every time point in the MCF-7 and T47D profiles. **(D)** Percentage of genes showing bimodal gene expression across 0, 1, 2, 3 and 4 time points among all genes in the MCF-7 and T47D profiles. Genes without any bimodal time point (red sector) are defined as unimodal genes; genes with at least 1 bimodal time point are defined as bimodal genes. **(E)** Average level (log2(FPKM+1)) of unimodal genes, bimodal genes, lower (mu1) and higher (mu2) components of bimodal genes across all MCF-7 and T47D single-cell profiles. **(F)** Density plots showing the expression distribution of known estrogen-responsive genes TFF1, GREB1, PCNA, and CA12 across single-cells in every time point. Line charts showing the average expression level (log_2_(FPKM+1)) in all single-cell profiles across time points; average level of lower (Mu1) and higher (Mu2) components for PCNA and CA12 are also shown. Open circle indicates significant alteration (P&0.05 by Mann-Whitney U test) compared to 0 h; closed circle indicates no significant alteration.

One of the major issues of a bimodal gene is that its averaged expression level value does not truly represent its two subpopulations. To address this, we compared the averaged expression level of bimodal and unimodal genes. As shown in Figure 3E, the average expression level of bimodal genes in general is significantly lower than the average expression level of unimodal genes. However, when we singled out the component 2 (higher-expressing subpopulation) of bimodal genes and compared it to unimodal genes, we found that the average expression level of component 2 (Mu2) is significantly higher than the average expression of the unimodal genes (Figure 3E). Thus, our findings suggest that highly expressed genes in a subpopulation of cells might be masked due to the averaging with the lower expressing subpopulation.

To explore this observation further, we examined the expression of 2 well-known E2-responsive genes (TFF1 and GREB1) with low bimodality and another 2 known E2-responsive genes (PCNA and CA12)^14^ with high bimodality (Figure 3F). We applied the Mann-Whitney U-test to examine whether their gene expression levels at 3, 6 and 12 hours after E2 stimulation changed significantly compared to that at 0 hour. As expected, the expression levels of both TFF1 and GREB1 changed significantly along the time course. For PCNA and CA12, we examined the expression level of all the cells in both components (Mu1 and Mu2). Intriguingly, Mu2 for PCNA changed significantly at 3 and 6 hours, but due to the averaging with Mu1 (which was not significantly changed), the mean expression level of all cells was not altered significantly. This example illustrates how a typical up-regulated gene in a subpopulation is masked in bulk. More interestingly, for CA12, even though both Mu1 and Mu2 were not significantly altered at 12 hours, the mean expression for all the cells was significantly changed. In this case, this is an example of Simpson’s paradox, a phenomenon in which the average of two groups displays an inverse trend compared to the two individual groups^15^. Similar averaging artifacts were also observed for known E2-responsive genes (PGR and CCND1)^14^ with high bimodality in T47D cells (Supplementary Figure 10). Taken together, our findings show that bimodal gene expression introduces different types of averaging artifacts and greatly affects the identification of DEGs.

### Pseudotemporal analysis identifies more differentially expressed genes with high bimodality

To resolve the averaging artifacts introduced by the bimodal expression of genes and at the same time obtain a better temporal resolution of the E2-stimulated transcriptional dynamics, we applied a recently developed computational method, “Monocle” ^11^, to reorder all the MCF-7 single-cell profiles along a pseudotemporal continuum (Figure 4A). Using this approach, we identified 1,102 differentially expressed genes (DEGs) based on the pseudotemporal continuum (Figure 4B). In comparison, 1,011 DEGs were obtained based on real-time points. We compared the differentially expressed genes from the two approaches and found 552 common genes (Figure 4B). We also found more bimodal DEGs based on pseudotime (813 genes) as compared to real-time (660 genes) (Figure 4C). Similar findings were also observed for T47D cells (Supplementary Figure 11). Among the pseudotime DEGs were well-known E2-responsive genes including TFF1, GREB1, PCNA, and CA12 (Figure 4D). Moreover, other E2-responsive genes such as CCNB2 and TOP2A^16^ were among the top bimodal DEGs (Supplementary Figure 12) that were not identified as DEGs based on real-time analysis, which highlights the power of the pseudotemporal method in solving the averaging artifact problem.

**Figure 4.**
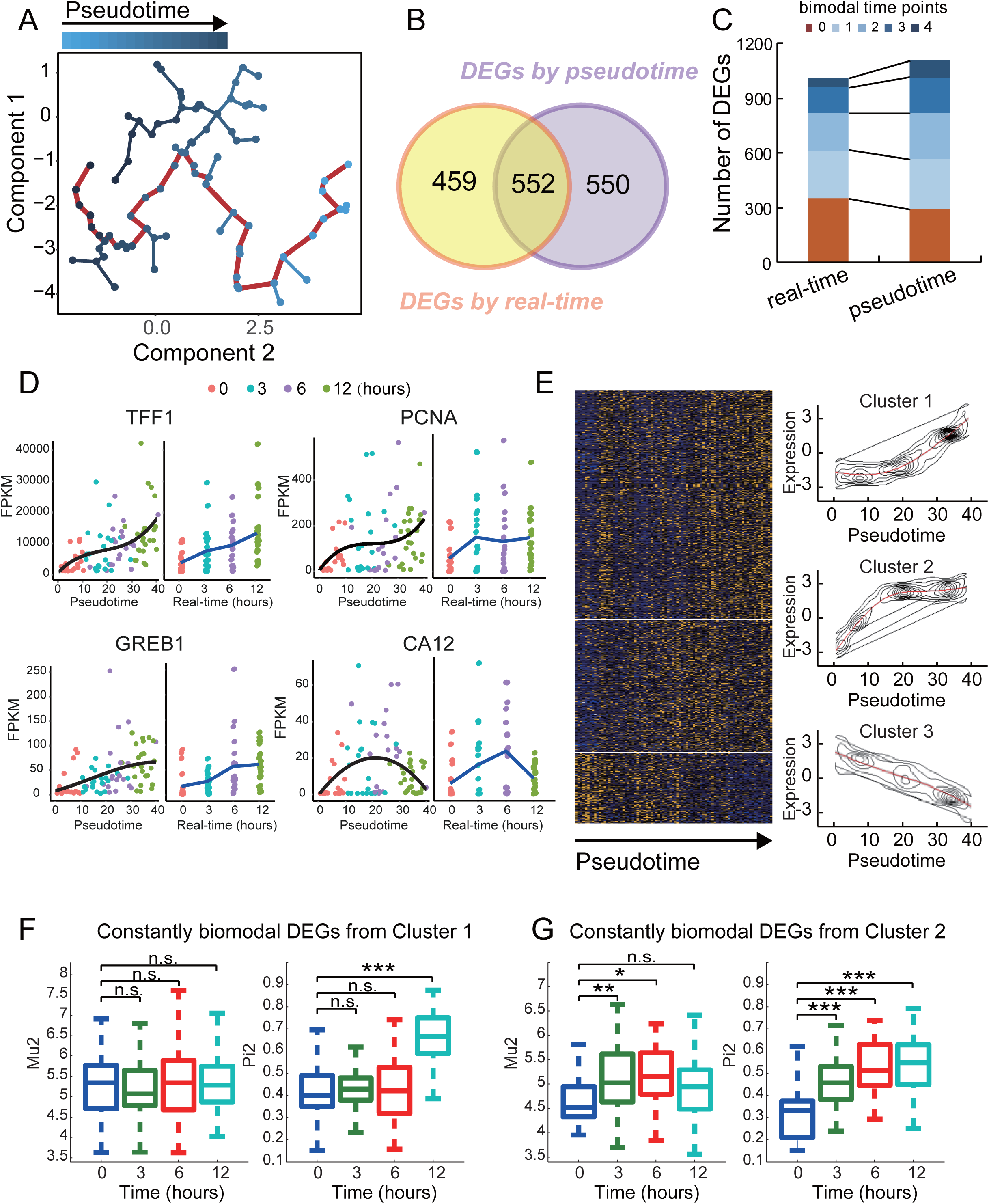
Pseudotime ordering of estrogen stimulated single-cell profiles by Monocle. **(A)** Single-cell profile (points) of MCF-7 cells in a two-dimensional independent component space. Lines connecting points represent edges of the minimal spanning tree (MST) constructed by Monocle. Solid red line indicates the main diameter of MST and provides the backbone of Monocle’s pseudotime ordering of the cells. **(B)** Number of DEGs identified based on real-time analysis and pseudotime analysis. **(C)** Number of DEGs with 0, 1, 2, 3 and 4 bimodal time points identified by Monocle based on real-time and pseudotime. **(D)** Expression of marker genes TFF1, GREB1, PCNA, and CA12 ordered by real-time and pseudotime. **(E)** Expression profiles for differentially expressed genes (DEGs) identified by Monocle based on pseudotime order. The DEGs were grouped into 3 clusters by k-means clustering. **(F)** Alterations of the component ratios and the average expression levels of constantly bimodal DEGs (bimodal in all the 4 time points) in Cluster 1 and 2. Pi2 is the ratio of the higher expressing subpopulation; Mu2 is the average expression level of the higher expressing subpopulation.

Next, we performed clustering analysis on the pseudotime DEGs and identified 3 distinct groups: Cluster 1 represents late upregulated genes; Cluster 2 represents early upregulated genes; and Cluster 3 represents repressed genes (Figure 4E). To characterize how bimodal DEGs change across the pseudotemporal continuum, we selected the constantly bimodal genes (63 in Clusters 1 and 23 in Cluster 2) for further analysis. Cluster 3 was not included as it has only 4 constantly bimodal genes. Surprisingly, we observed Pi2 values, but not Mu2 values significantly changed at 12 hours (Figure 4F and G). This finding suggests that most up-regulated bimodal DEGs are “switched-on” to a similar expression level in more cells rather than elevated to a higher expression level in all the cells.

### Estrogen stimulates metabolic reprogramming that favors biosynthesis and cell proliferation but reduces degradation of estrogen

To gain functional insights into the DEGs identified by pseudotemporal analysis, we performed pathway enrichment analysis. As shown in Figure 5A, upregulated DEGs are enriched in pathways that favor cell proliferation including cell cycle, DNA repair, and anabolic pathways. In contrast, downregulated DEGs are enriched in the xenobiotic and glutathione metabolism pathways (Figure 5A). These two pathways are associated with the degradation of estrogen-derived byproducts and ROS clearance^17^. Thus, the downregulation of these two pathways could potentially lead to an accumulation of genotoxic estrogen metabolites and an elevation of reactive oxygen species (ROS), which are known to promote tumorigenesis^18^. Taken together, our findings indicate that E2 stimulation induces a metabolic switch that favors biosynthesis and cell proliferation, but impairs estrogen catabolism and ROS clearance.

**Figure 5.**
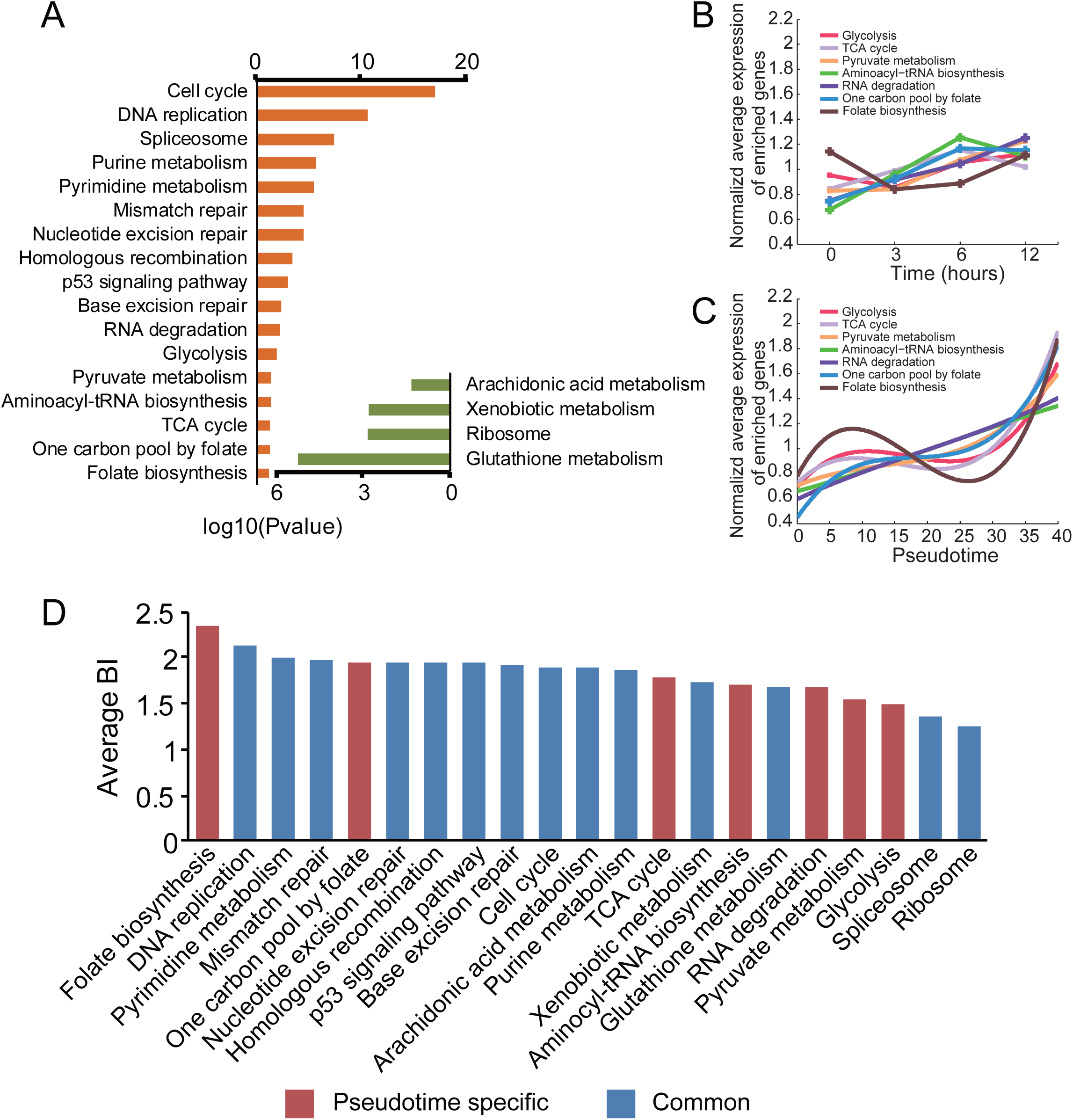
Estrogen-stimulated metabolic reprogramming in MCF-7 cells. **(A)** KEGG pathways significantly enriched among the DEGs based on pseudotime. **(B)** Expression trend of up-regulated pathways based on real time. **(C)** Expression trend of up-regulated pathways based on pseudotime. Expression level of a pathway was calculated by averaging the normalized levels of DEGs within the pathway. **(D)** Average BI values of the up-regulated pathways. Average BI value of a pathway is calculated by averaging the BI values of all DEGs in all time points in the pathway. Pathways identified by pseudotime only are indicated in red.

Among the above enriched pathways, glycolysis, TCA cycle, aminoacyl-tRNA biosynthesis, one-carbon metabolism, and folate biosynthesis pathways were identified only by pseudotime analysis but not by real-time analysis. By examining the expression trend of these pathways based on real-time points (Figure 5B) and pseudotime (Figure 5C), we found that in general the pseudotime analysis showed much greater alterations in these pathways. In addition, we found genes in the one-carbon metabolism and folate biosynthesis pathways having very high BI values; therefore, these two pathways were missed by real-time analysis possibly due to the averaging artifacts of gene bimodality (Figure 5D). To the best of our knowledge, the regulation of these two pathways by estrogen in breast cancer cells has not been reported in previous time-series studies of bulk MCF-7 cells using either gene expression microarray^4^, RNA-sequencing^14^ or proteomics^19^ technologies.

### Estrogen stimulates reprogramming of folate-mediated one-carbon metabolism through the mitochondrial folate pathway

Since our analysis revealed folate and one-carbon metabolism are novel estrogen-responsive pathways, we decided to investigate these two pathways in further detail. Folate metabolism is a pivotal process for one-carbon metabolism and thus they are often linked together and called folate-mediated one-carbon metabolism^20^. Folate metabolism can fuel one-carbon metabolism via either the mitochondrial pathway or the cytosolic pathway^21^. As shown in Figure 6A, E2 upregulated the expression of key enzymes in the mitochondrial pathway including SHMT2, MTHFD2, and MTHFD1, but not SHMT1, a key enzyme in the cytosolic pathway (data not shown). Hence, our results suggest folate-mediated one-carbon metabolism is upregulated preferentially via the mitochondrial folate pathway in response to E2 stimulation. E2 also stimulated other important enzymes that are associated with this pathway such as DHFR, GGH, TYMS, and MAT2B (Figure 6A), which likely contributes to the upregulation of nucleotide synthesis, DNA repair, DNA methylation and polyamine synthesis pathways. Interestingly, most of the above genes exhibited a bimodal expression patterns (Figure 6B, Supplementary Figure 13, left panel). Moreover, the expression trend based on pseudotime changed more significantly than those based on real-time points (Figure 6B, Supplementary Figure 13, right panel). Taken together, our observations for the folate-mediated one carbon metabolism pathway is consistent with our earlier conclusion that the increase in expression of bimodal genes is due to the ratio of higher-expressing subpopulation and not by the average expression level in the total cells.

**Figure 6.**
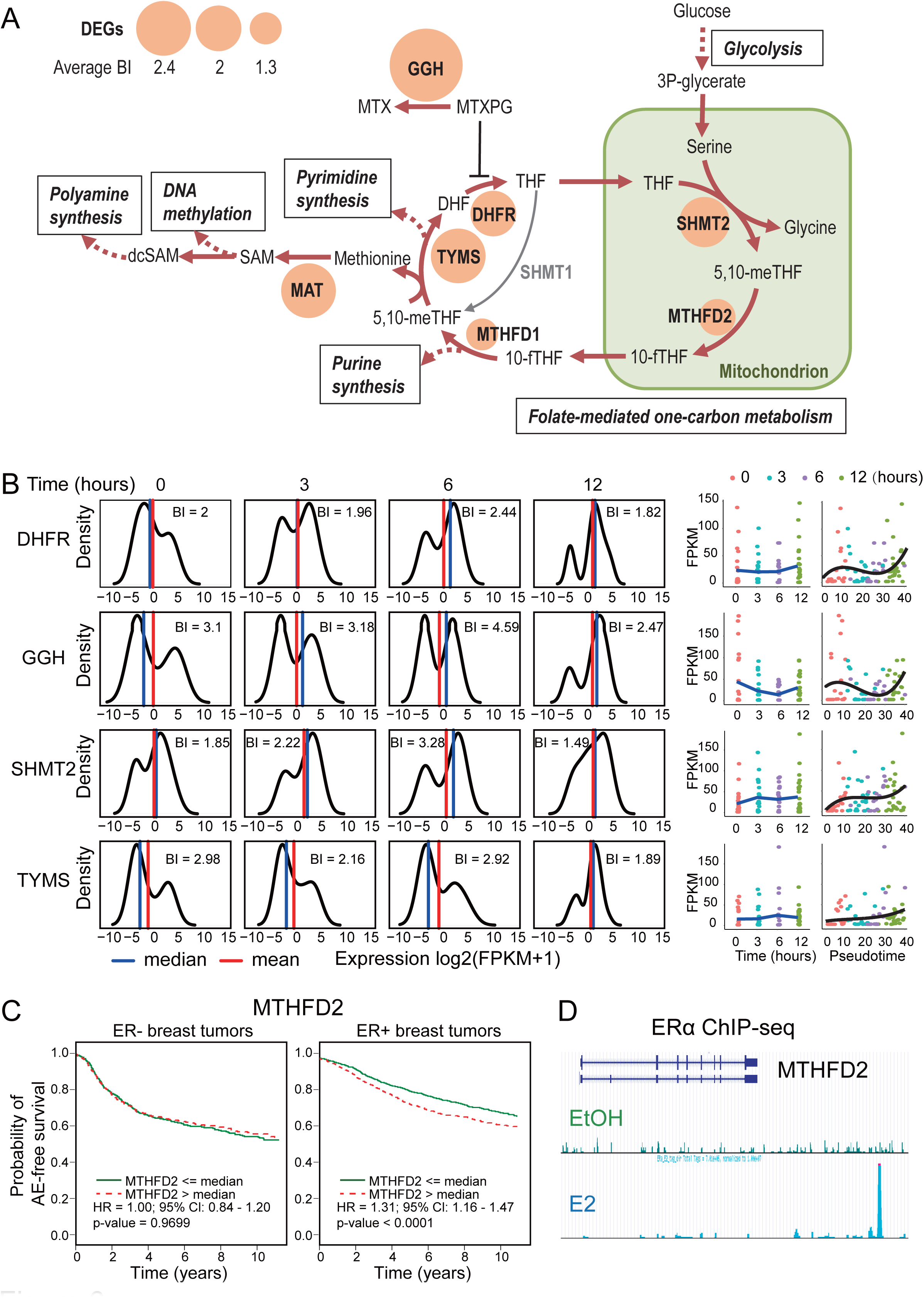
Estrogen stimulates reprogramming of one-carbon metabolism through the mitochondrial folate pathway. **(A)** Schematic presentation showing significantly altered key regulators in the folate-mediated one-carbon metabolism pathway and their average BI values. THF, tetrahydrofolate; 5,10-meTHF, 5,10-methylenetetrahydrofolate; 10-fTHF, 10-formyltetrahydrofolate; DHF, dihydrofolate; SAM, S-adenosylmethionine; dcSAM, decarboxylated S-adenosylmethionine; MTX, methotrexate; MTXPG, methotrexate polyglutamates. **(B)** DEGs with high BI values in the folate biosynthesis and one-carbon metabolism pathways. Density plots show the expression level distribution in every time point. Lines charts show expression trend of the genes based on real time and pseudotime. **(C)** Kaplan-Meier survival analysis of MTHFD2 for ERα-positive and negative breast cancer patients. **(D)** ERα ChIP-seq data in MCF-7 cells shows binding sites near the MTHFD2 gene body.

Next, we asked whether any of these genes play an important role in ERα-positive breast tumor progression. For this, we performed prognostic analysis on the key enzymes in the folate-mediated one-carbon metabolism pathway using bc-GenExMiner 4.0^22^, a webtool based on a collection of 36 breast cancer cohort studies. Our results showed that these genes could potentially serve as good prognostic markers for ERα-positive but not ERα-negative breast cancer patients (Figure 6C, Supplementary Figure 14). Moreover, we asked whether any of these genes are direct targets of ERα. Using our previous ERα ChIP-seq data in MCF-7^23^, we found an ERα binding site located 12 kb upstream of the transcription start site (TSS) of MTHFD2, suggesting that this gene is likely a direct target of ERα (Figure 6D). MTHFD2 and the mitochondrial folate pathway have been shown to be important for various solid tumors^24^, but its dependence on estrogen regulation in ERα-positive breast tumors has not been reported yet. Taken together, our pseudotemporal analysis revealed that estrogen reprograms one-carbon metabolism through the mitochondrial folate pathway.

### Estrogen stimulation augments mitochondrial folate pathway to coordinately fuel polyamine and de novo purine synthesis

When we examined the metabolic pathways associated with one-carbon metabolism in further detail, we found both the polyamine and de novo purine synthesis pathways are also coordinately regulated by estrogen stimulation (Figure 7A). Specifically, the upstream enzymes and regulators of these pathways including MTHFD2, PPAT, and AZIN1 are early E2 upregulated genes (Figure 7A, Supplementary Figure 15). In contrast, several of the enzymes in the downstream part of the pathways such as MTHFD1, PAICS, and ODC1 are late E2 upregulated genes (Figure 7A, Supplementary Figure 15). The early responsive genes AZIN1 and PPAT are also bimodal DEGs discovered by pseudotemporal analysis but not by real-time point analysis (Supplementary Figure 15). Integrative analysis with ERα ChIP-seq data^25^ showed both genes are associated with ERα binding sites in MCF-7 cells and ERα-positive breast tumors (Figure 7B and C, left panel). In survival analysis, we also found these genes are significant prognostic markers for ERα-positive but not ERα-negative breast cancer patients (Figure 7B and C, right panel). Taken together, these findings suggest that AZIN1 and PPAT are important ERα targets in ERα-positive breast tumors.

**Figure 7.**
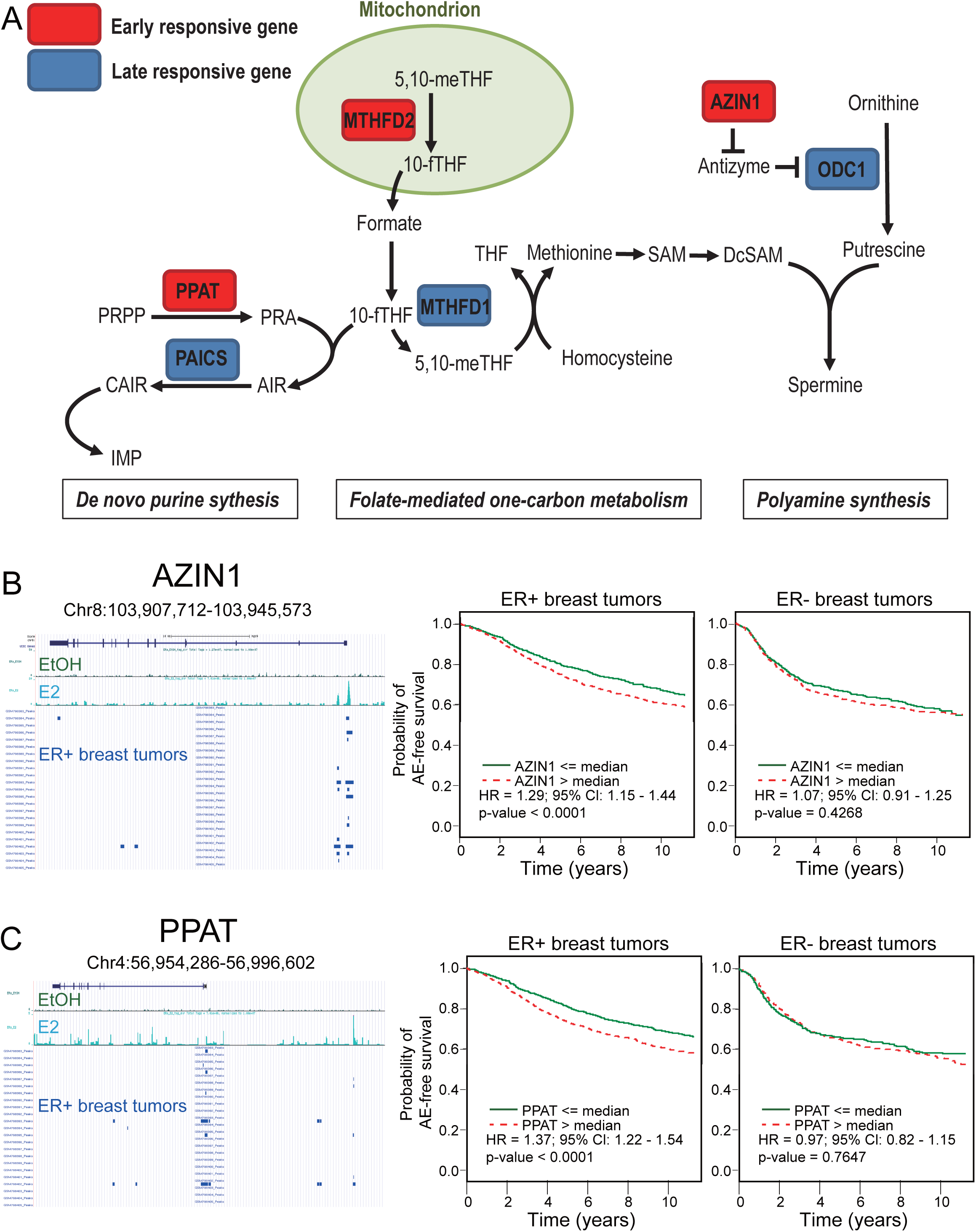
Estrogen stimulation coordinately reprograms one-carbon metabolism to fuel purine and polyamine synthesis. **(A)** Schematic presentation showing estrogen stimulation coordinately regulates one-carbon metabolism, de novo purine synthesis and polyaminesynthesis. PRPP, phosphoribosyl pyrophosphate; PRA, phosphoribosylamine; AIR, aminoimidazole ribotide; CAIR, carboxyaminoimidazole ribotide. **(B)** ChIP-seq data showing ERα binding sites near the AZIN1 gene in MCF-7 cells and in clinical tumor samples. Kaplan-Meier survival analysis of AZIN1 for ERα-positive and negative breast cancer patients. **(C)** ChIP-seq data showing ERα binding sites near PPAT gene bodies in MCF-7 cells and in clinical tumor samples. Kaplan-Meier survival analysis of PPAT for ERα-positive and negative breast cancer patients.

To examine whether AZIN1 and PPAT have important functional roles in ERα-positive breast cancer, we conducted dsiRNA knockdown assays for AZIN1 and PPAT in MCF-7 cells. MTT cell proliferation assays showed that knockdown of both genes dramatically reduced MCF-7 cell proliferation (Figure 8A and B). In addition, we performed sub-G1 assay and observed a significant increase in cell death in MCF-7 cells after gene knockdown (Figure 8C and D). Finally, to determine the reason for the cell death, we performed Annexin V/propidium iodide assay and showed that gene knockdown induced strong cell apoptosis in MCF-7 cells. Therefore, AZIN1 and PPAT are essential genes for ERα-positive breast cancer cell proliferation and survival.

**Figure 8.**
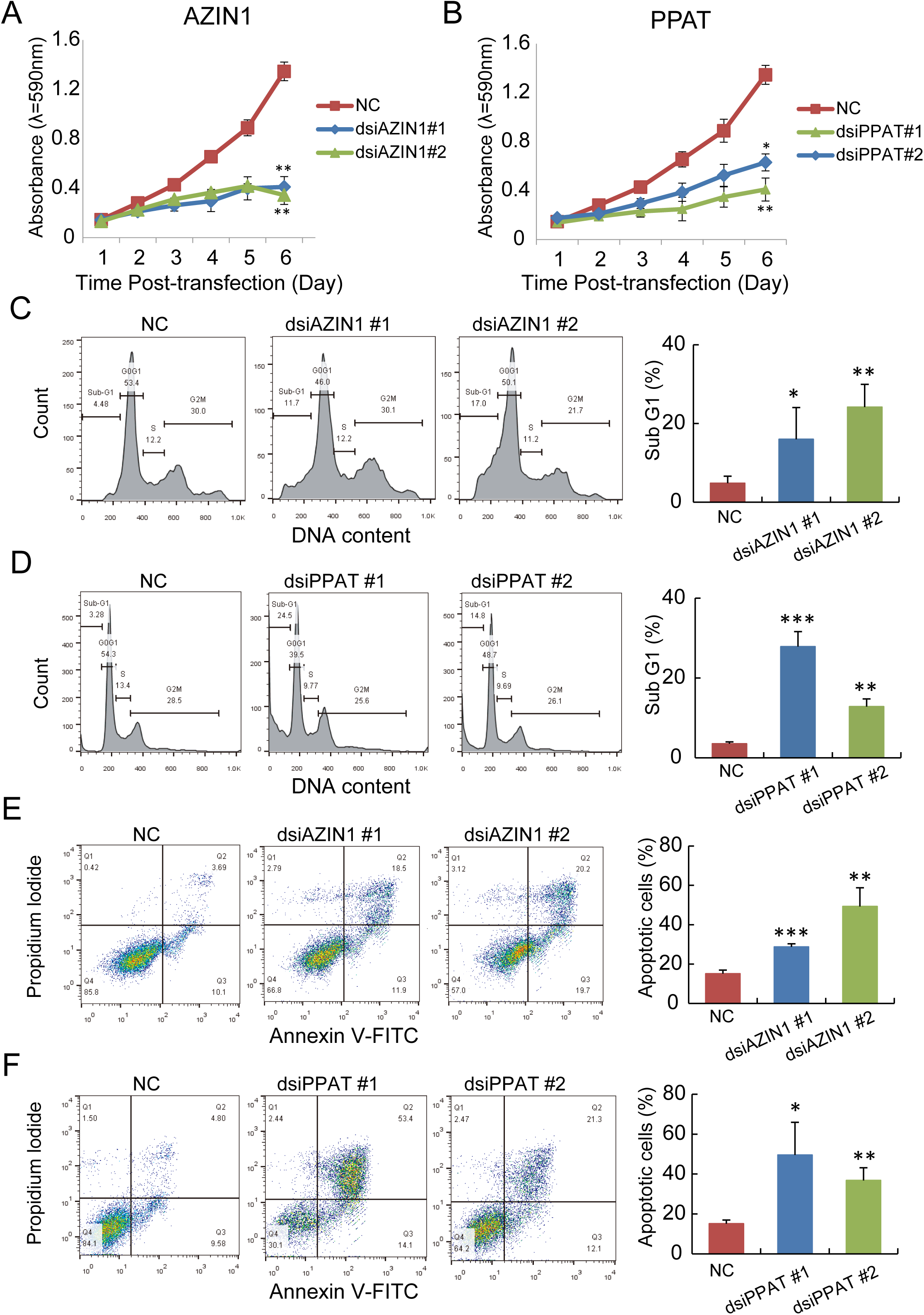
AZIN1 and PPAT are essential for breast cancer cell proliferation and survival. **(A)** Knockdown of AZIN1 with dsiRNAs reduces MCF-7 cell proliferation. **(B)** Knockdown of PPAT with dsiRNAs reduces MCF-7 cell proliferation. **(C)** Knockdown of AZIN1 with dsiRNAs increases MCF-7 sub-G1 cell fraction. **(D)** Knockdown of PPAT with dsiRNAs increases MCF-7 sub-G1 cell fraction. **(E)** Knockdown of AZIN1 with dsiRNAs increases MCF-7 cell apoptosis. **(F)** Knockdown of PPAT with dsiRNAs increases MCF-7 cell apoptosis. Error bars represent SD. *, P & 0.05; **, P & 0.01; ***, P & 0.001.

In conclusion, we have revealed a novel metabolic switch process orchestrated by E2 regulation in breast cancer cells. Upon E2 stimulation, ERα directly upregulates expression of MTHFD2, AZIN1 and PPAT to coordinately fuel polyamine and purine synthesis via the mitochondrial folate pathway.

## Discussion

In this study, we performed single-cell RNA-seq analysis to profile the transcriptional landscape of E2 time-course treatment on ERα-positive breast cancer cell lines. We used the Fluidigm C1 platform to facilitate high-throughput automated single-cell capture and cDNA preparation of nearly 200 single-cell libraries from MCF-7 and T47D cells. To our knowledge, this is the first study to characterize the cellular transcriptional response to estrogen stimulation at the single-cell level.

As expected, we observed a greater variation in gene expression between individual cell transcriptomes compared to bulk population of cells. However, the intercellular correlation of our datasets (0.65 < r < 0.80) was higher than previously published datasets derived from in vivo cells (0.29 < r < 0.62)^9^. We believe this difference could be in part due to the more uniformed in vitro cell culture environment in our experiment. As a validation of our scRNA-seq results, we used ddPCR and smRNA-FISH, two independent techniques that are capable of accurate absolute quantification. Our results suggest the cell-to-cell heterogeneity that we observed in our scRNA-seq is real and not due to technical noise because of sequencing minute amounts of material.

A recent single-cell study examining the response of lipopolysaccharide (LPS) stimulation on mouse bone marrow-derived dendritic cells showed that most LPS-responsive genes are highly variable and more than 75% of the highly variable genes exhibit a bimodal expression pattern^9^. These findings are consistent with our observations in breast cancer cells in which we also identified a large proportion of E2-responsive genes with bimodal expression. Specifically, we found more than two thirds of the genes showed bimodal expression in at least one time point. More importantly, the gene expression bimodality in many instances produced averaging artifacts. First, it masked the higher expression subpopulation by averaging it with the lower expression subpopulation. Second, the expression trend of the subpopulation was masked. In some extreme examples, Simpson’s paradox happened, that is, the expression trend of the total cells and the two subpopulations were inverse to each other. From our work, we not only observed the gene expression bimodality but also explained some of the possible artifacts that caused by it when using conventional methods for DEG analysis.

To circumvent this problem, we employed the Monocle algorithm which order single-cell profiles along an artificial pseudotemoral continuum^11^. Each single-cell profile was considered as a pseudotime point on the continuum and thus substantially enhanced the temporal resolution. The DEGs that were identified based on real-time points and pseudotime were quite different (Figure 4B). In particular, there was significantly more bimodal genes identified by pseudotime as compared to real-time (Figure 4C). Moreover, analysis of the constantly bimodal DEGs showed that these highly bimodal genes were altered significantly by the ratios of subpopulations instead of the averaged expression level of the subpopulations (Figure 4F and G). These results highlight a new model of transcriptional regulation by nuclear receptors in response to hormone stimulation. Unlike unimodal genes that change in expression levels, the activated levels of these bimodal genes are tightly restricted at a narrow range in individual cells, but the frequencies of “switch-on” increase showing a higher ratio of the switched-on subpopulation. Monocle successfully identified more highly bimodal responsive genes, probably due to the higher temporal resolution by a continuous arrangement of single-cell profiles instead of discrete grouping by a few experimental time points. Therefore, we have demonstrated that the single-cell trajectory analysis method is a good solution for dissecting the averaging artifacts produced by gene expression bimodality.

From our single-cell analysis, we identified folate synthesis and one-carbon metabolism as novel estrogen-regulated metabolic pathways that are enriched for bimodal genes (Figure 6B). Folate metabolism can fuel one-carbon metabolism via either the mitochondrial folate pathway or the cytosolic folate pathway^21^. Interestingly, we found that estrogen stimulation did not upregulate the cytosolic folate pathway but instead selectively activated the mitochondrial folate pathway (Figure 6A). Moreover, we identified MTHFD2 as a novel ERα target and an early responsive regulator for the reprogramming. MTHFD2 and the mitochondrial folate pathway have recently been highlighted as new markers for metabolic reprogramming in a wide spectrum of solid tumors^24^. Thus, the link between ERα regulation and MTHFD2 overexpression in our finding underscores an important event for the establishment of metabolic reprogramming in ERα-positive breast tumors. On the other hand, glycolysis and TCA cycle pathways, which are known estrogen-regulated pathways in breast cancer^26^, were also identified by pseudotemporal analysis but not by real-time point analysis. These pathways are enriched for unimodal genes; hence, increased temporal resolution not only helped to identify highly bimodal responsive genes but also aided in the identification of unimodal responsive genes with insignificant changes between discrete time points.

Our single-cell analysis also showed the polyamine and de novo purine synthesis pathways are coordinately upregulated by estrogen with the mitochondrial folate pathway. In these pathways, we identified the bimodal genes, AZIN1 and PPAT, as early E2 responsive genes and novel ER□ targets. AZIN1 is an inhibitor of antizyme that induces degradation of ODC1 and interferes with the uptake of external polyamine^27^, while PPAT is the first-step rate-limiting enzyme for de novo purine synthesis^28^. Other important enzymes downstream of AZIN1 and PPAT, such as ODC1^27^ and PAICS^28^ are among the late E2 responsive genes. Hence, the transcriptional dynamics illustrated by our pseudotemporal analysis is consistent with the flow of the metabolic pathways (Figure 7A). Finally, we were able to validate both AZIN1 and PPAT are essential for MCF-7 proliferation and migration (Figure 8). Although the mitochondrial folate pathway is adopted by various tumors^24^, recent studies showed that it can be compensated by the cytosolic folate pathway if it is disrupted^21^. Therefore, targeting the polyamine and de novo purine synthesis pathways may be a better choice than targeting the folate mitochondrial pathway.

In summary, the dynamic heterogeneity and metabolic switch of cellular response to E2 stimulation revealed from our work may serve as a potential strategy for cells to eventually acquire resistance to hormone therapy. Thus, we believe further single-cell transcriptome analysis of additional breast cancer cell lines in long-term anti-estrogen and aromatase-inhibition treatments will bring invaluable insight for understanding how therapy-resistant cells emerge from a given population of cancer cells.

## Materials and Methods

### Cell Lines

MCF-7 and T47D ERα-positive human breast cancer cell lines were obtained from ATCC and maintained in DMEM and RPMI 1640 media (Gibco), respectively. Both media were supplemented with 10% fetal bovine serum (FBS) (Gibco) and grown in a 37°C incubator with 5% CO2.

### E2 stimulation and scRNA-seq library generation

Three days prior to E2 stimulation, MCF-7 and T47D cells were transferred into phenol-red free DMEM/F12 and RPMI media (Gibco), respectively. Both media were supplemented with 10% charcoal/dextran-stripped FBS (Hyclone). Cells were then stained with CellTracker Green CMFDA or Orange CMTMR (Molecular Probes) 24 h before drug treatment. Stained cells were treated with either ethanol (vehicle) or E2 at 4 time points (0 and 3 h: Green, 6 and 12 h: Orange), before being dissociated into single-cell suspension for single-cell and bulk population analyses.

Single-cell cDNA samples were obtained using the Fluidigm C1 Single-Cell Auto Prep System. Briefly, 125,000 cells from two time points (one labeled red and the other labeled green) were mixed together in 1 ml of media and then loaded into a Fluidigm C1 Integrated Fluidic Chip (IFC). The IFC was loaded into the C1 Single-Cell Auto Prep System for cell capture. After cell capture, the IFC was imaged using an automated inverted fluorescence microscope to identify the captured cells based on their fluorescence. Then cell lysis and cDNA synthesis processes were continued on the chip. Finally, single-cell cDNA samples were harvested as per manufacturer’s instructions.

Cells from each time point were also used in bulk population cDNA generation. For these cells, RNA was extracted using the RNeasy Mini Kit (QIAGEN) and then reverse transcribed using the SMARTer Ultra Low RNA Kit (Clontech) according to the manufacturer’s protocol.

Microscope images of the single-cells were examined to exclude wells with multiple cells or cells with multiple staining. Single-cell and bulk population samples were barcoded and pooled together for library preparation using the Nextera XT Sample Prep Kit (Illumina). The pooled libraries were quantified through quantitative PCR using the KAPA Library Quantification Kit (KAPA Biosystems), followed by sequencing on the Illumina platform (2 x 76 bp for the MCF-7 libraries or 2 x 101 bp for the T47D libraries). The scRNA-seq data from this work have been deposited in the Gene Expression Omnibus (GEO) under the accession number GSE107864.

### scRNA-seq data processing

All RNA-seq reads were first trimmed to 75 bp. Next, the adapter sequences were trimmed. Reads were mapped as paired-end sequencing reads with the Tophat package (v.2.0.6) against UCSC hg19 and the gene annotation reference Gencode V18. The FPKM (Fragments Per Kilobase of exon model per Million mapped fragments) expression values were calculated from the BAM file generated by Tophat using the Cufflink package (v.2.0.2). Genes with FPKM greater than one in at least one sample were retained for further analysis. Sequencing quality was evaluated using the FastQC and RSeQC packages to assess the single-cell libraries on the total sequenced reads, uniquely mapped reads, 5’ or 3’ coverage bias, as well as the percentage of intronic bases and intergenic bases. A sample was deemed biased if the coverage coefficient of the median gene body position was less than 0.8 (Supplementary Figure 2). Single-cell libraries with a low sequencing depth of fewer than 1 million reads or exhibiting 5’ or 3’ coverage bias were filtered out. In total, 84 MCF-7 and 78 T47D single-cell samples (Supplementary Table 1) were used for downstream analyses. We performed unsupervised clustering on all the bulk population and single-cell samples based on Pearson correlation coefficients of the global expression profiles between every two samples.

### Gene expression bimodality analysis

A total of 5,975 coding genes were assessed for bimodality expression using the R package, SIBER^13^, which is a bioinformatic approach for identifying bimodal expressed genes from RNA-seq data. The SIBER algorithm fits the single-cell gene expression distribution into 2 log-normal distributions (component 1 and 2) and calculates the mean values (Mu1 and Mu2) and the ratios (Pi1 and Pi2) for these two components. A bimodal index (BI) value was calculated to assess the potential bimodality for each gene by the following equation:

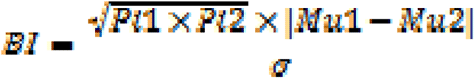

We defined a gene with a BI value > 1.8 (generated from 1,000 times random repeats and top 5% confident degree) in at least one time point as a bimodally expressed gene.

### Mann-Whitney rank sum test

Genes with FPKM < 1 in more than median number of single-cell samples were filtered. We compared samples from the 3, 6, and 12 h time points with those from the 0 h time point. We performed Mann-Whitney rank sum test (U-test) on the genes and considered P < 0.05 as significant DEGs.

### Pseudotemporal analysis

Pseudotemporal analysis of the single-cell gene expression data was performed using the R package, Monocle^11^. The Monocle algorithm implemented dimension reduction by independent component analysis (ICA) prior to cell ordering. The artificial temporal curve was generated by minimal spanning trees (MST) that connect cells along the longest possible path involving as many cells as possible. DEGs across the pseudotime axis were identified using a generalized additive model (GAM). Testing for differential expression was performed with an approximate χ2 likelihood ratio test.

KEGG pathway enrichment analysis of the DEGs was performed using the DAVID database^29^. Prognostic analysis of selected genes in ERα-positive and negative tumors was performed using bc-GenExMiner 4.0 with 5,861 clinical cases from a collection of 36 previous breast cancer cohort studies^22^. ERα ChIP-seq data from breast tumor tissues were retrieved from GSE32222 in GEO DataSets^25^.

### Single-cell droplet digital PCR

Single-cell cDNA samples were diluted by 100 times and 1 μL was used for detecting the estrogen-responsive genes, ESR1, TFF1, and GREB1, and the house-keeping gene, ACTB. Duplexed droplet digital PCR (ddPCR) was performed according to the manufacturer’s protocol and the results were reported as copies of mRNA per μL sample. In brief, 1 μL of DNA template was added to 20 μL of ddPCR Supermix for Probes (Bio-Rad) and primer/probe mixtures of ESR1 (FAM probe) + ACTB (HEX probe) or TFF1 (FAM probe) + GREB1 (HEX probe). The 20 μL samples and 70 μL of Droplet Generation Oil for Probes (Bio-Rad) were used for droplet generation. Droplets were then thermal cycled with the following conditions: 5 min at 95℃, 40 cycles of 94℃ for 30 sec, 55℃ for 1 min followed by 98℃ for 10 min (ramp rate 2C/sec). Samples were then transferred to a QX200 Droplet Reader (Bio-Rad) for fluorescent measurement of FAM and HEX signals. Gating was performed on the basis of negative controls and the copies of mRNA were calculated for each sample using QuantaSoft software (Bio-Rad).

### Single-molecule RNA fluorescent in situ hybridization

Multiplexed single-molecule RNA fluorescent in situ hybridization (smRNA-FISH) was performed using RNAscope probes (ACDBio) according to the manufacturer’s protocol. MCF-7 and T47D cells were treated with E2 treatment at 0, 3, 6 and 12 hr. TFF1, GREB1, and ESR1 mRNAs were all probed at the same time. At least 200 cells per condition were counted for the mRNA copies for each gene. Each experiment was repeated 3 independent times.

### Cell proliferation assays

To assess the effects of AZIN1 and PPAT depletion on MCF-7 cell proliferation, cells were first transfected with dsiRNAs targeting AZIN1, PPAT, or a non-targeting (NC) control. 24 h after transfection, cells were then seeded into 96-well plates at 3 × 10^3^ cells/well. Cells were incubated with 5 mg/mL sterilized MTT (Life Technologies) in PBS for 3 h at 37°C. The plate was then incubated for a further 30 min at 37°C. The absorbance was measured at an OD of 590 nm using a plate reader. Cell proliferation was monitored every day over a period of 6 days. Experiments were performed in triplicates and repeated 3 independent times.

### Cell apoptosis assays

Cell apoptosis assay was performed using the Dead Cell Apoptosis Kit (Invitrogen) according to the manufacturer’s instruction. In brief, MCF-7 cells were transfected with dsiRNAs targeting AZIN1, PPAT or a non-targeting (NC) control. After 72 h of transfection, the cells were harvested, centrifuged at 1,200 rpm for 5 min at 4°C, and then resuspended in binding buffer containing Annexin V-FITC and propidium iodide. The cells were incubated for 15 min in the dark at room temperature. After staining, the cells were mixed with 400 μl binding buffer and immediately analyzed by flow cytometry. Experiments were repeated 3 independent times.

### Statistical Analysis

The results are represented as mean ± s.d. The statistical significance of differences was determined by paired, two-tailed Student’s t-Test. A p-value of <0.05 was considered to be statistically significant.

## Acknowledgement

This work was supported by the Macau Science and Technology Development Fund (FDCT/023/2014/A1 and FDCT102/2015/A3), a University of Macau Multi-Year Research Grant (MYRG2015-00196-FHS), and a University of Macau Start-up Research Grant (SRG2014-0000-FHS) to EC.

## Supplementary Information

**Supplementary Table 1.** Number of single-cell libraries generated

**Supplementary Table 2.** Primer sets and probes used in droplet digital PCR

**Supplementary Table 3.** DsiRNAs used for knockdown assays

**Supplementary Figure 1.** Box-plots of the distribution of unique mapping reads for the MCF-7 and T47D single-cell profiles.

**Supplementary Figure 2.** Coverage of the unique mapping reads across gene bodies for the MCF-7 and T47D single-cell profiles.

**Supplementary Figure 3.** Distribution of correlation coefficients for all MCF-7 and T47D single-cell pairs showing high degree of variability (R=0.65-0.75).

**Supplementary Figure 4.** Variation of gene expression in the MCF-7 and T47D bulk and single-cell profiles. Variation (x-axis) is the median absolute deviation of the FPKM (MAD/M). For each gene, the MAD/M is plotted against the log2-transformed median FPKM value to visualize how variation changes with overall transcript abundance.

**Supplementary Figure 5.** Quantification of TFF1 and GREB1 mRNA copies in cDNA samples by duplexed droplet digital PCR. **(A)** Number of droplets generated for the serial dilution of MCF-7 cDNA samples by BioRad QX200 Droplet Generator. Graph showing all the droplet numbers are above 16,000. **(B)** Quantification of TFF1-positive and GREB1-positive droplets simultaneously in cDNA samples by ddPCR. **(C)** Quantification of TFF1-positive droplets in serially diluted cDNA samples. **(D)** Quantification of GREB1-positive droplets in serial diluted cDNA samples. **(E)** Counts for TFF1 mRNA copies in 10X series dilution cDNA samples. **(F)** Counts for GREB1 mRNA copies in 10X serial dilution cDNA samples.

**Supplementary Figure 6.** Quantification of ESR1 and ACTB mRNA copies in cDNA samples bt duplexed droplet digital PCR. **(A)** Number of droplets generated for the serial dilution MCF-7 cDNA samples by BioRad QX200 Droplet Generator. Graph showing all the droplet numbers are above 16,000. **(B)** Quantification of ESR1-positive and ACTB-positive droplets simultaneously in cDNA samples by ddPCR. **(C)** Quantification of ESR1-positive droplets in series dilution cDNA samples. **(D)** Quantification of ACTB-positive droplets in serially diluted cDNA samples. **(E)** Counts for ESR1 mRNA copies in 10X serially diluted cDNA samples. **(F)** Counts for ACTB mRNA copies in 10X serially diluted cDNA samples.

**Supplementary Figure 7.** Quantification of mRNA copies of TFF1, GREB1, ESR1, and ACTB in single-cell cDNA samples by ddPCR.

**Supplementary Figure 8.** Correlation between mRNA copies quantified by ddPCR and FPKM value quantified by RNA-seq of TFF1, GREB1, ESR1, and ACTB in single-cell cDNA samples.

**Supplementary Figure 9. (A)** Violin plots presenting RNA-seq expression values (log2(FPKM+1)) of estrogen-responsive marker genes ESR1, GREB1, and PGR for T47D single-cells in time-course E2 treatment. **(B)** Violin plots presenting smRNA-FISH values (log2(Count+1)) of the marker genes for T47D single-cells in an independent time-course E2 treatment. 200 cells were counted for each time point. **(C)** Single-molecule RNA-FISH to quantified mRNA copies of the gene markers in T47D single-cells under time-course estrogen treatment. Nuclei are counter stained by DAPI.

**Supplementary Figure 10.** Examples of averaging artifacts due to gene bimodality in T47D single-cells. Density plots show the expression level distribution of known estrogen-responsive genes PGR and CCND1 across single-cells in every time point. Line charts show average expression levels, Mu1 and Mu2 in all single-cell profiles across time points. Open circle indicates significant alteration (P<0.05 by Mann-Whitney U test) compared to 0 h; close circle indicates no significant alteration.

**Supplementary Figure 11. (A)** Single-cell profile (points) of T47D in a two-dimensional independent component space. Lines connecting points represent edges of the minimal spanning tree (MST) constructed by Monocle. Solid red line indicates the main diameter of MST and provides the backbone of Monocle’s pseudotime ordering of the cells. **(B)** Number of DEGs identified based on real time points and pseudotime. **(C)** Percentage of DEGs with 0, 1, 2, 3 and 4 bimodal time points identified based on real time and pseudotime. **(D)** Expression profile for differentially expressed genes (DEGs) identified by Monocle based on pseudotime order. The DEGs are further grouped into 3 clusters based k-means clustering.

**Supplementary Figure 12.** Known ERα direct targets that have high BI values and are identified by pseudotime only.

**Supplementary Figure 13.** Other DEGs in the folate-mediated one-carbon metabolism pathway (correlated to Figure 6B).

**Supplementary Figure 14.** Kaplan-Meier survival analysis and Forest plots for DEGs in the folate-mediated one-carbon pathway (TYMS, SHMT2, DHFR, GGH, MTHFD2, MTHFD1) for ERα-positive and negative breast cancer patients (correlated to Figure 6C).

**Supplementary Figure 15.** DEGs in the polyamine and de novo purine synthesis pathways.

## References

1 DeSantis, C., Ma, J., Bryan, L. & Jemal, A. Breast cancer statistics, 2013. CA Cancer J Clin 64, 52–62, doi:10.3322/caac.21203 (2014).

2 Cheung, E. & Kraus, W. L. Genomic analyses of hormone signaling and gene regulation. Annu Rev Physiol 72, 191–218, doi:10.1146/annurev-physiol-021909-135840 (2010).

3 Liu, M. H. & Cheung, E. Estrogen receptor-mediated long-range chromatin interactions and transcription in breast cancer. Mol Cell Endocrinol 382, 624–632, doi:10.1016/j.mce.2013.09.019 (2014).

4 Frasor, J. et al. Profiling of estrogen up- and down-regulated gene expression in human breast cancer cells: insights into gene networks and pathways underlying estrogenic control of proliferation and cell phenotype. Endocrinology 144, 4562–4574, doi:10.1210/en.2003-0567 (2003).

5 Metivier, R. et al. Estrogen receptor-alpha directs ordered, cyclical, and combinatorial recruitment of cofactors on a natural target promoter. Cell 115, 751–763 (2003).

6 Battich, N., Stoeger, T. & Pelkmans, L. Control of Transcript Variability in Single Mammalian Cells. Cell 163, 1596–1610, doi:10.1016/j.cell.2015.11.018 (2015).

7 Harper, M. A. et al. Phenotype sequencing: identifying the genes that cause a phenotype directly from pooled sequencing of independent mutants. PLoS One 6, e16517, doi:10.1371/journal.pone.0016517 (2011).

8 Leng, N. et al. Oscope identifies oscillatory genes in unsynchronized single-cell RNA-seq experiments. Nat Methods 12, 947–950, doi:10.1038/nmeth.3549 (2015).

9 Shalek, A. K. et al. Single-cell transcriptomics reveals bimodality in expression and splicing in immune cells. Nature 498, 236–240, doi:10.1038/nature12172 (2013).

10 Shalek, A. K. et al. Single-cell RNA-seq reveals dynamic paracrine control of cellular variation. Nature 510, 363–369, doi:10.1038/nature13437 (2014).

11 Trapnell, C. et al. The dynamics and regulators of cell fate decisions are revealed by pseudotemporal ordering of single cells. Nat Biotechnol 32, 381–386, doi:10.1038/nbt.2859 (2014).

12 Durruthy-Durruthy, R. & Heller, S. Applications for single cell trajectory analysis in inner ear development and regeneration. Cell Tissue Res 361, 49–57, doi:10.1007/s00441-014-2079-2 (2015).

13 Tong, P., Chen, Y., Su, X. & Coombes, K. R. SIBER: systematic identification of bimodally expressed genes using RNAseq data. Bioinformatics 29, 605–613, doi:10.1093/bioinformatics/bts713 (2013).

14 Yamaga, R. et al. RNA sequencing of MCF-7 breast cancer cells identifies novel estrogen-responsive genes with functional estrogen receptor-binding sites in the vicinity of their transcription start sites. Horm Cancer 4, 222–232, doi:10.1007/s12672-013-0140-3 (2013).

15 Simpson, E. H. The interpretation of interaction in contingency tables. J R Stat Soc Series B Stat Methodol 13, 238–241 (1951).

16 Walker, G. et al. Estrogen-regulated gene expression predicts response to endocrine therapy in patients with ovarian cancer. Gynecol Oncol 106, 461–468, doi:10.1016/j.ygyno.2007.05.009 (2007).

17 Santen, R. J., Yue, W. & Wang, J. P. Estrogen metabolites and breast cancer. Steroids 99, 61–66, doi:10.1016/j.steroids.2014.08.003 (2015).

18 Yager, J. D. Mechanisms of estrogen carcinogenesis: The role of E2/E1-quinone metabolites suggests new approaches to preventive intervention‐‐A review. Steroids 99, 56–60, doi:10.1016/j.steroids.2014.08.006 (2015).

19 Drabovich, A. P., Pavlou, M. P., Dimitromanolakis, A. & Diamandis, E. P. Quantitative analysis of energy metabolic pathways in MCF-7 breast cancer cells by selected reaction monitoring assay. Mol Cell Proteomics 11, 422–434, doi:10.1074/mcp.M111.015214 (2012).

20 Ducker, G. S. & Rabinowitz, J. D. One-Carbon Metabolism in Health and Disease. Cell Metab 25, 27–42, doi:10.1016/j.cmet.2016.08.009 (2017).

21 Ducker, G. S. et al. Reversal of Cytosolic One-Carbon Flux Compensates for Loss of the Mitochondrial Folate Pathway. Cell Metab 24, 640–641, doi:10.1016/j.cmet.2016.09.011 (2016).

22 Jezequel, P. et al. bc-GenExMiner: an easy-to-use online platform for gene prognostic analyses in breast cancer. Breast Cancer Res Treat 131, 765–775, doi:10.1007/s10549-011-1457-7 (2012).

23 Joseph, R. et al. Integrative model of genomic factors for determining binding site selection by estrogen receptor-alpha. Mol Syst Biol 6, 456 (2010).

24 Nilsson, R. et al. Metabolic enzyme expression highlights a key role for MTHFD2 and the mitochondrial folate pathway in cancer. Nat Commun 5, 3128, doi:10.1038/ncomms4128 (2014).

25 Ross-Innes, C. S. et al. Differential oestrogen receptor binding is associated with clinical outcome in breast cancer. Nature 481, 389–393, doi:10.1038/nature10730 (2012).

26 Forbes, N. S., Meadows, A. L., Clark, D. S. & Blanch, H. W. Estradiol stimulates the biosynthetic pathways of breast cancer cells: detection by metabolic flux analysis. Metab Eng 8, 639–652, doi:10.1016/j.ymben.2006.06.005 (2006).

27 Kahana, C. Antizyme and antizyme inhibitor, a regulatory tango. Cell Mol Life Sci 66, 2479–2488, doi:10.1007/s00018-009-0033-3 (2009).

28 Pedley, A. M. & Benkovic, S. J. A New View into the Regulation of Purine Metabolism: The Purinosome. Trends Biochem Sci 42, 141–154, doi:10.1016/j.tibs.2016.09.009 (2017).

29 Huang da, W., Sherman, B. T. & Lempicki, R. A. Systematic and integrative analysis of large gene lists using DAVID bioinformatics resources. Nat Protoc 4, 44–57, doi:10.1038/nprot.2008.211 (2009).

